# Ancestral proteins trace the emergence of substrate specificity and oligomerization within bacterial DEDDy dinucleases

**DOI:** 10.1101/2025.03.30.646021

**Authors:** Sofia Mortensen, Andy Burnim, Keith Dufault-Thompson, Alexandra E. Lipka, Xiaofang Jiang, Holger Sondermann

**Affiliations:** CSSB Centre for Structural Systems Biology, Deutsches Elektronen-Synchrotron DESY, Notkestr. 85, 22607 Hamburg, Germany; National Center for Biotechnology Information, National Library of Medicine, National Institutes of Health, 8600 Rockville Pike, Bethesda, MD 20894, USA; Christian-Albrechts-University, 24118 Kiel, Germany

## Abstract

Nucleases play a crucial role in bacterial physiology, influencing processes such as DNA repair, genome maintenance, and host-pathogen interactions. We recently identified a class of nucleases, diDNases, which are encoded on mobile genetic elements and homologous to the house-keeping nanoRNase C (NrnC). Despite their shared structural fold, diDNases and NrnC orthologs exhibit differences. DiDNases form dimers and preferably cleave DNA dinucleotides, whereas NrnC homologs assemble into octamers that do not discriminate between RNA or DNA dinucleotides. Here, we investigate the evolutionary divergence of these enzymes using ancestral sequence reconstruction. Our results show that both diDNases and NrnC orthologs originated from a dimeric ancestor with intermediate substrate preferences. Structural analyses of ancestral and extant dinucleases provide a molecular rational for how gradual changes in conformation gave rise to substrate preferences, oligomeric state, and catalytic efficiency of these related, yet distinct enzyme clades. These findings provide insights into how small structural modifications enable large-scale changes in molecular assembly and functional specialization harnessing a conserved protein fold. In addition, the preference of the early ancestors for DNA dinucleotides and preservation of this activity in all extant enzymes strongly argues for a biological function of DNA dinucleotides.

## Introduction

Mobile genetic elements (MGE) play important roles in bacterial evolution and have great influence on the ecology and dynamics of bacterial populations. MGEs come with fitness cost for a host but, at the same time, may provide benefits in form of offensive and defensive functions, antibiotic resistance, and pathogenicity genes. Due to the close co-evolution of organisms and MGEs, genes from an organism’s core genome may become mobilized while genes within MGEs may be incorporated into the host’s core genome, becoming domesticated in that lineage (*1*).

We recently discovered a distinct group of enzymes, dubbed diDNases, preferentially encoded in MGEs of Actinomycetes and Clostridia that are homologs of a house-keeping nanoRNase C (NrnC) found in Alphaproteobacteria, Cyanobacteria, and Spirochaetia (*2*). DiDNases and NrnC orthologs have a lot in common: they are both members of the DEDDy subfamily of nucleases and utilize a strict two Mg^2+^ ion catalysis to cleave phosphodiester bonds. Additionally, both nucleases share a common fold. However, there are significant differences between diDNases and NrnC orthologs with regard to their substrate preferences and physiological role. As their name indicates, diDNases are highly specific towards 5’-phosphorylated, single-stranded DNA of two nucleotides in length, deoxydinucleotides, whereas NrnC orthologs cleave both DNA and RNA dinucleotides with equal efficiency (*2*). Even though NrnC as well as the functionally analogous oligoribonuclease (Orn) have been implicated in essential processes of catalyzing the final degradation step of RNA and RNA-based second messengers, the relevance of their activity on DNA remains enigmatic (*3*–*7*). In contrast, based on their genomic neighborhood, diDNases are predicted to play a role in DNA mobility or phage defense (*2*). Overall, the high sequence and structural similarity of NrnC orthologs and diDNases stands in striking contrast to their differential activities and physiological roles.

Yet another difference between NrnC and diDNase is in their quaternary structure. Both enzymes dimerize using similar interfaces, but NrnC orthologs assemble further into octamers comprising four dimeric subunits, while diDNases do not show signs of higher oligomerization beyond a constitutive dimeric form (*2*). The importance of the higher-order oligomeric arrangement of NrnC is still unclear.

Structural comparison of the enzyme-substrate complexes of diDNase and NrnC revealed that functional differences of the nucleases could not be explained only by single amino acid residue substitutions, but rely on large-scale conformational alterations (*2*). In order to understand these factors and to explore how striking variation in biochemical properties arose within highly similar structural folds, we employed bioinformatics and experimental approaches to resurrect protein ancestors of extant diDNases and NrnC orthologs. We present here a functional analysis of the reconstructed ancestral proteins revealing that evolution of diDNases and NrnC orthologs started from an ancestor with intermediate properties that bifurcated towards more or less DNA preference. We also present a crystal structure of an ancestor, which allowed us to pinpoint structural alterations that led to the emergence of the specific activities of extant diDNases and NrnC orthologs. We provide evidence that octamerization of NrnC is likely not a random evolutionary event and is linked to overall increased enzymatic activity.

## Results

### Resurrection of diDNase/NrnC ancestral proteins

DiDNases and NrnC orthologs share significant sequence similarity and adopt a common fold; on the other hand, they have striking distinctions in substrate preference and quaternary structure (2), which cannot readily be explained by localized sequence differences between the two DEDDy protein families. To develop a fundamental understanding of how the differential properties in the enzymes evolved in a shared fold, we chose a bioinformatics approach relying on ancestor reconstruction.

We constructed a comprehensive phylogenic tree of NrnC orthologs, diDNases, and closely related homologous proteins and mapped the proteins with known substrate preferences on the tree based on previous reports (*2, 3*) (Fig. 1A). This phylogenetic reconstruction suggests that the divergence between the diDNase and NrnC clades occurred relatively early in the evolution of the enzyme family, with the NrnC clade then further splitting into distinct sub-clades associated with the Cyanobacteria, Spirochaetia, and Alphaproteobacteria phyla (Fig. 1A). The bootstrap values for the split between diDNases and the NrnC groups are well supported (bootstrap > 90). However, the support values for the splits between NrnC clades were relatively low, making it difficult to determine the directionality of evolution within the NrnC family. We then utilized ancestral sequence reconstruction to predict the likely ancestral sequences of enzymes at key branching points in the tree. We identified five key nodes based on the tree topology that represented common ancestors and divergence points between clades of interest (Fig. 1B). We designed the sequence of common diDNase/NrnC ancestor A0, the ancestor of the diDNase clade – A1, the ancestor of the entire NrnC clade – A2, and ancestors of the sub-clades on NrnC – A3, A4, and A5. Sequence alignment of the ancestors showed that the characteristic DEDDy dinuclease signatures at EXO I, EXO II, EXO III motifs, and at the C-terminus emerged later than A0, A1, and A2 branching points (Fig. S1A).

**Figure 1.**
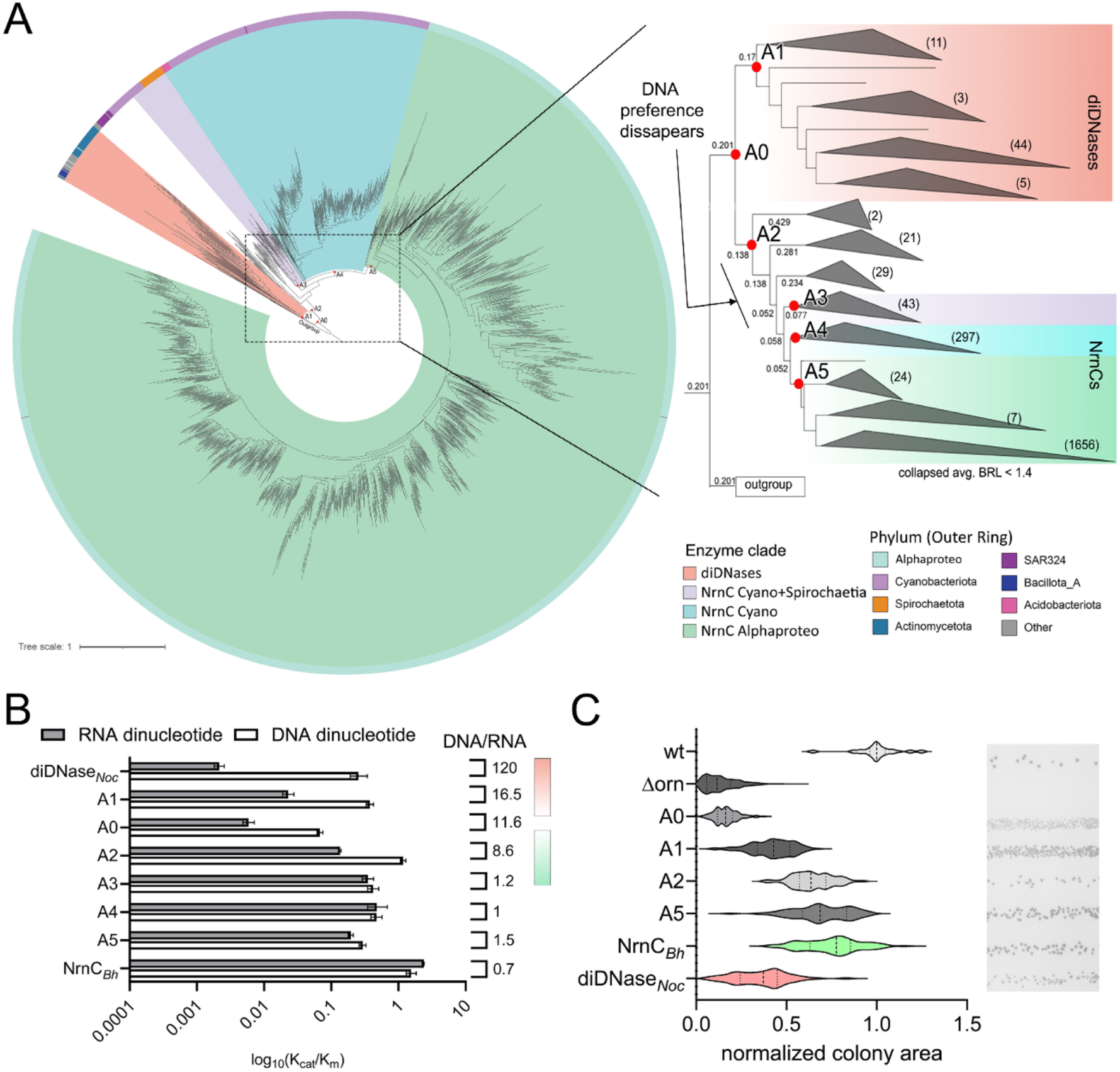
Ancestral protein reconstruction of diDNases and NrnC orthologs. **A**. Phylogenic tree with diDNase and NrnC clades, manually rooted using homologous outgroup sequences. Taxonomic assignments are mapped on the outer ring. Branch length between important nodes are shown in the close up view of the ancestral nodes between extant diDNase and NrnC clades. Leaves of the tree with average branch length < 1.4 were collapsed and the number of leaves in each collapsed clade are stated in parenthesis. Ancestral nodes chosen for characterization are marked with a red dot. **B**. Catalytic efficiency of ancestral protein toward DNA and RNA dinucleotides, dp(2AP)G and p(2AP)G, respectively. Values for kcat /Km (×10^6^ M^−1^ s^−1^) are plotted on a log10 scale. The error bars show 95% confidence interval of the model fitting. The efficiency values of diDNase from *Nocardioides* (diDNase_*Noc*_) and NrnC from *Bartonella* (NrnC_*Bh*_) are plotted as references. Gradient color bar represent specification with regards for DNA preference going into two different directions. **C**. Complementation of the small-colony phenotype of *P. aeruginosa Δorn* using the indicated ancestral proteins and controls of NrnC_*Bh*_ and diDNase_*Noc*_. Violin plots show colony sizes normalized to the average size of the wild-type *P. aeruginosa* colonies from the same plate. A representative image of a drip assay plate used for scoring is shown in the right panel.

In order to understand the evolution of substrate specificity of diDNases with their strong preference for DNA on one hand and NrnC proteins that do not discriminate between RNA and DNA dinucleotides as substrates on the other hand, we recombinantly expressed the resurrected ancestral sequences, purified the proteins and measured their enzymatic activities towards RNA and DNA dinucleotides (Fig. 1C, S1B, Table S1). All the ancestors were enzymatically active and showed significant nuclease activity (Fig. S1B, Table S1). However, both the common diDNase/NrnC ancestor A0 and diDNase ancestor A1 displayed somewhat lower levels of activity towards both substrates as compared to extant NrnC orthologs and diDNases (*2*). Ancestor A0 had the lowest overall level of activity among all measured ancestral and extant proteins: 66-442 times lower activity for RNA and 7-23 times lower for DNA. In case of ancestor A1, activity compared to that of extant enzymes was 16-104 times and 1-4 times lower for RNA and DNA, respectively (Table S1). These data indicate that the dinuclease function of the proteins was optimized after the emergence of the common ancestors A0, A1, and A2. Nevertheless, while the ancestors A0, A1, and A2 cleaved both DNA and RNA nucleotides, they demonstrated higher catalytic efficiency towards DNA than RNA dinucleotide (Fig. 1C, S1B, Table S1). The DNA preference gradually increased in the diDNase lineage and decreased in NrnC lineage. In the NrnC lineage, the preference for DNA substrates was lost completely after the A2 ancestor leading to the extant NrnC proteins, with ancestors A3 to A5 displaying roughly equal activity on RNA and DNA dinucleotides (Fig. 1B-C, S1B, Table S1). In the lineage leading to diDNases, strong DNA preference emerged after the A1 ancestor. Additionally, ancestor A1 already had an activity level toward DNA substrates that was comparable to that of extant diDNases. To summarize, we showed that the evolution of substrate specificity started from enzymes with slight preference for DNA dinucleotides and went two ways, toward stronger DNA preference in the diDNase clade and a lack of preference in the NrnC clade.

In order to examine whether the *in vitro* enzyme kinetics correlate with dinuclease activities of the ancestor *in vivo*, we tested the ability of ancestors A0, A1, A2, and A5 to complement the small-colony phenotype of *Pseudomonas aeruginosa Δorn* (Fig. 1C). Orn possesses comparable activity on RNA and DNA dinucleotides and a complete rescue of the *orn* deletion like depends on this feature (*2*). Hence, the level of small-colony phenotype complementation indicates whether a protein, when expressed in the mutant strain, acts as a RNA/DNA non-discriminating dinuclease or an enzyme with DNA preference, *e*.*g*. a diDNase, which would result in an incomplete phenotype rescue (*2*–*4*). Ancestor A0 provided almost no complementation consistent with its low overall nuclease function and slight preference for DNA observed *in vitro*. Ancestor A1 could only rescue to an average level of 0.5 relative to the average colony size wild-type *P. aeruginosa*. A similar complementation potential was observed for an extant diDNase (Fig. 1C) (*2*). Ancestors A2 and A5 showed complementation levels identical to a non-discriminating NrnC. Together, these results suggest that Orn can only be replaced by enzymes that cleave both RNA and DNA dinucleotides efficiently; an enzyme with a strong preference for DNA dinucleotides provides only a partial rescue.

**Figure S1.**
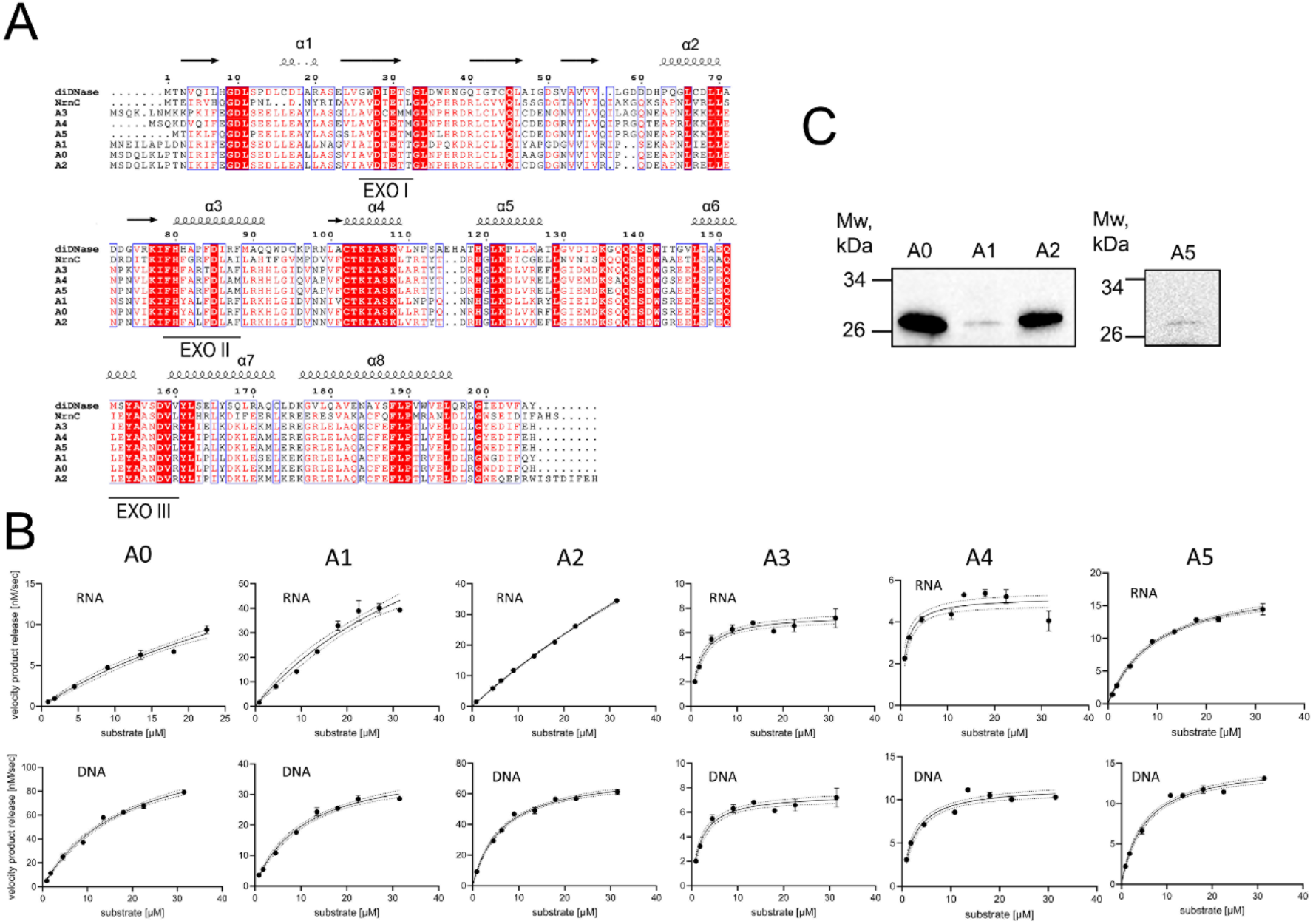
Sequence and functional analysis of the ancestral proteins. **A**. Sequence alignment of ancestor protein at nodes of interest and *Bartonella henselae* NrnC (NrnC_*Bh*_) and *Nocardioides* diDNase (diDNase_*Noc*_). **B**. Enzymatic activity of ancestral proteins towards RNA (p(2AP)G) and DNA (dp(2AP)G) substrates. Data were fitted with a Michaelis-Menten model. **C**. Western blot detection of the indicated StrepII-tagged ancestral proteins as expressed from an inducible plasmid in the *P. aeruginosa Δorn* strain. Expression of NrnC_*Bh*_ and diDNase_*Noc*_ were established in (2).

**Table S1.** Enzyme kinetics parameters of the ancestral proteins towards DNA and RNA dinucleotide substrates.

|  | A0 |  | A1 |  | A2 |  | A3 |  | A4 |  | A5 |  |
| --- | --- | --- | --- | --- | --- | --- | --- | --- | --- | --- | --- | --- |
|  | DNA | RNA | DNA | RNA | DNA | RNA | DNA | RNA | DNA | RNA | DNA | RNA |
| [E], nM | 100 | 100 | 10 | 100 | 10 | 10 | 10 | 10 | 10 | 10 | 10 | 10 |
| V<sub>max</sub>, nM/sec | 126.40 | 27.31 | 40.47 | 108.00 | 74.18 | 178.00 | 14.09 | 7.50 | 11.56 | 5.16 | 15.04 | 19.05 |
| K<sub>m</sub>, μM | 18.67 | 46.67 | 10.78 | 47.33 | 6.35 | 131.60 | 3.37 | 2.14 | 2.47 | 1.09 | 5.16 | 9.76 |
| k<sub>cat</sub>, sec<sup>-1</sup> | 1.26 | 0.27 | 4.05 | 1.08 | 7.42 | 17.80 | 1.41 | 0.75 | 1.16 | 0.52 | 1.50 | 1.91 |
| k<sub>cat</sub>/K<sub>m</sub>, ×10<sup>6</sup> M<sup>-1</sup>s<sup>-1</sup> | 0.0677 | 0.0059 | 0.3754 | 0.0228 | 1.1689 | 0.1353 | 0.4180 | 0.3508 | 0.4682 | 0.4750 | 0.2916 | 0.1952 |
| DNA/RNA | 11.57 |  | 16.45 |  | 8.64 |  | 1.19 |  | 0.99 |  | 1.49 |  |

### Structural basis for the emergence of substrate preferences in diDNases and NrnC orthologs

In extant diDNases and NrnC orthologs, substrate preference was shown to correlate with a combination of large-scale structural features, namely dimer conformation, and more localized features, mainly implicating amino acid residues at positions 80, 87, and 205 (per *Bartonella henselae* NrnC, NrnC_*Bh*_, numbering). However, it was unclear to what extent the large-scale differences contribute to substrate restrictions. Also, the nature of the residues 80, 87, and 205 alone could not fully explain apparent substrate preferences, since swap mutations did not show effects on specificity or led to impaired overall enzyme activity (*2*). Therefore, we utilized the resurrected proteins to investigate whether the emergence of large-scale structural features, in particular in the dimeric protein core, between ancestors and extant enzymes tracks the divergence in substrate preferences of these enzymes. To this end, we determined a crystal structure of an ancestor with an intermediate level of DNA preference and high level of DNase activity, the ancestor A1, at 1.45 Å resolution. Since it was shown before that the presence of substrates in active sites of NrnC and diDNase does not lead to any major conformational changes (*2, 3*), we compared the protein entities of structures of diDNase_*Noc*_-dpGG (PDB ID 9F7G), NrnC_*Bh*_-pGG (PDB ID 7MPL), and ancestor A1 by aligning them on the catalytic motifs EXOI and EXOII of protomer A (Fig. 2A). Consistent with previous observations (*2*), the alignment showed variations in position of the C-terminus of protomer B in the active site of protomer A, and consequently variations in space at the active site for an RNA substrate, in all three structures. Additionally, helices α4 and α7, surrounding the C-terminus, are situated differently, which is best described by the angles between well aligned reference helix α3 and helix α4 of the protomer A and between helix α3 of A and α7 of B (Fig. 2A). The angle between helices α3 and α4 ranges from 157.6° in NrnC to 147.8° in diDNase; in ancestor A1 the corresponding angle measures 149°. The angle between α3 and α7 ranges from 162.2° in NrnC and to 126.4° in diDNase, with the two helices forming a 155.1° angle in ancestor A1. In the diDNase, the C-terminus together with helices α4, and α7 come closer to the reference helix α3 and the active site of a protomer A compared to their positions in NrnC. Similar to the angle measurements, these structural elements adopt intermediate positions in the ancestor A1 (Fig. 2A). This result together with our enzyme activity data suggest that alterations within just 2.5 Å of how deep the C-terminus of one protomer is placed into the active site of the other protomer in the dimeric unit appear crucial in tuning preference for DNA up or down during diDNase and NrnC evolution. These alterations were brought about by the changes in the conformation of helices α4 and α7.

**Figure 2.**
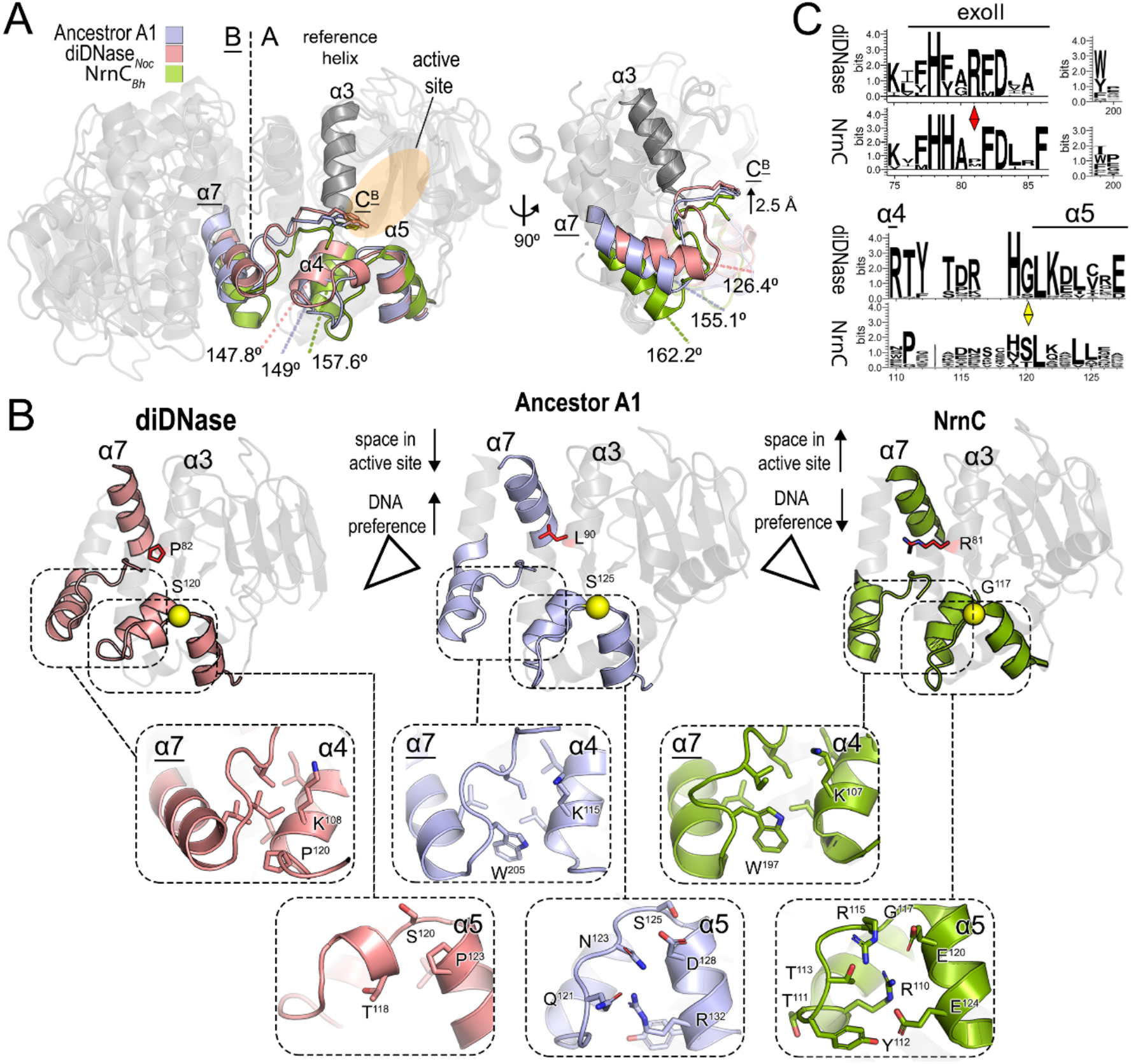
Structural basis for the evolution of substrate preferences in diDNases and NrnC orthologs. **A**. Comparison of structures of extant diDNase_*Noc*_-dpGG (PDB ID 9F7G), NrnC_*Bh*_-pGG (PDB ID 7MPL), and ancestor A1. The structures were aligned on the catalytic motifs EXOI and EXOII. The well aligned helix α3 in protomer A was chosen as a reference to measure the angles between helix α3 and helices α4 and α7, implicated in large conformational changes leading to a differential ability of active site to accept RNA as substrates. **B**. Detailed views of the structural differences, which cause alteration in positions of helices α4 and α7, in extant diDNase, NrnC, and ancestral A1 protein. The hinge position occupied by either glycine (G) or serine (S) is shown as a sphere, the position of the R wedge is shown in red. **C**. Sequence logos of the regions identified by structural analysis as defining the amount of space in the active site and, hence, DNA preference.

To rationalize these changes, we examined the factors influencing the conformations of the helices and their interactions with other elements (Fig. 2B). The ability of helix α4 to form a wide angle with helix α3 in NrnC can be attributed to a conserved glycine, G^117^, acting as a hinge at the end of the loop connecting helices α4 and α5, which provides more flexibility than the serine at the same position in diDNase and ancestor A1 (Fig. 2B-C). Another NrnC-specific element, which wedges between helix α7 and the C-terminus and prevents the C-terminus from reaching deeper into the active site, is residue R^81^ (Fig. 2B). Notably, R^81^ is highly conserved as a part of the EXOII motif in NrnC (Fig. 2C). In the ancestor A1 and diDNase_*Noc*_, the analogous positions are occupied by smaller and uncharged residues, leucine and proline, respectively. Additionally, in NrnC and ancestor A1, unlike in diDNase, there are several polar interactions between helix α5 and the α4-α5 loop, which hold helix α4 relatively further away from α3 (Fig 2B). The residues participating in those interactions are conserved in NrnC orthologs (Fig. 2C). The distance between helices α4 and α7 in both NrnC and ancestor A1 is slightly larger compared to that in diDNases, likely due to the presence of a conserved, bulky residue W^197^ in the hydrophobic interface between helices α4 and α7 of NrnC and ancestor A1. Taken together, ancestral structure A1 possesses a mixture of NrnC- and diDNase-specific elements (Fig. 2B), ultimately leading to its intermediate level of DNA preference.

### Co-emergence of substrate promiscuity and octamer assembly in NrnC orthologs

In addition to distinct substrate preferences, NrnC orthologs and diDNases differ in their oligomeric states (2). DiDNases form stable dimers, while NrnC orthologs analyzed so far exist as tetramers of the analogous dimeric units, forming an overall octameric assembly (2–4). In order to understand the appearance of the octamerization propensity during NrnC evolution and functional specialization we mapped the quaternary structure phenotypes of the extant enzymes and the computed ancestors on the phylogenic tree we inferred (Fig. 3A). The experimental oligomerization assignments of twelve extant enzymes (*Rhodococcus, Nocardioides, Clostridium, Blastococcus, Cellulosimicrobium, Bartonella, Brucella, Agrobacterium, Inquilinus, Leptospira, Okeania, Synechococcus*) were determined previously (2–4). Here, we experimentally characterized oligomeric states of additional NrnC orthologs (Fig. S2A) as well as of the ancestors by size-exclusion chromatography-coupled multi-angle light scattering (SEC-MALS) (Fig. 3B). The NrnC orthologs invariably formed octamers in solution under the experimental conditions, a feature shared with ancestors A3, A4, and A5. In contrast, ancestors A0, A1, and A2 formed stable dimers under the same conditions, akin to diDNases.

**Figure 3.**
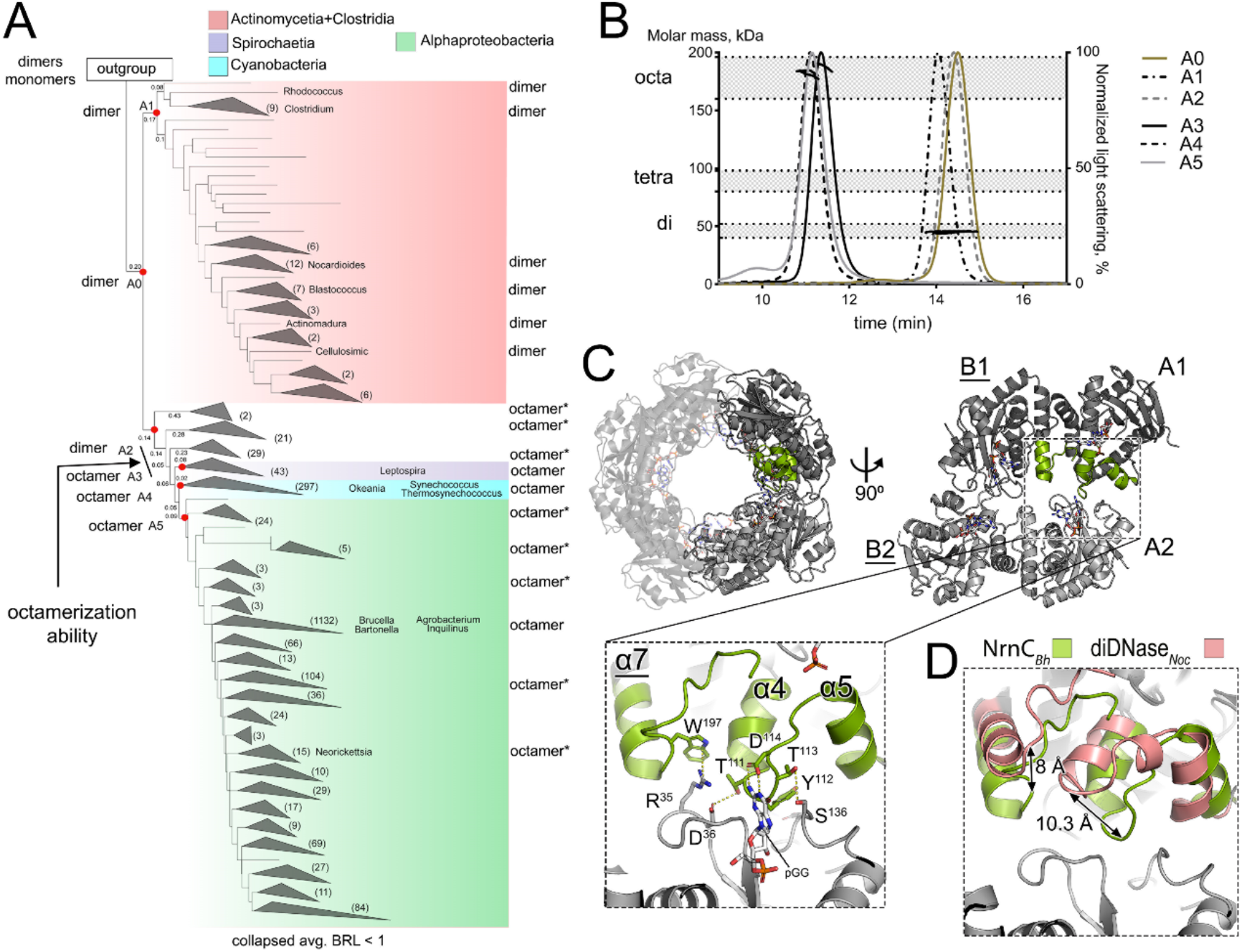
Emergence of octamerization in NrnC. **A**. Mapping of the oligomeric state phenotype onto the phylogenic tree and ancestral nodes. Branch length between important nodes are shown. Leaves of the tree with average branch length < 1.0 were collapsed and the number of leaves in each collapsed clade are stated in parenthesis. Ancestral nodes chosen for characterization are marked with a red dot. Oligomeric states determined either experimentally or by using AlphaFold3 predictions (*) are indicated. **B**. Oligomeric state of ancestral proteins determined by SEC-MALS. The normalized light scattering signal (chromatogram lines, right axis) and molar mass values (dots across peaks, left axis) were plotted against the SEC elution volume. Shaded areas show the theoretical molar mass ranges for dimeric, tetrameric and octameric assemblies based on the primary sequence of the proteins. **C**. Structural analysis of the interfaces between dimeric subunits inside a NrnC octamer (PDB ID 7MPL) pinpoint the interactions holding the units together. **D**. Close-up view of the NrnC octameric interface area with a diDNase structure (PDB ID 9F7G) overlaid on it showing the major differences in the position of the interacting helices α4, α5 and α7, and the loop α4-α5.

To cover parts of the phylogenic tree lacking experimental oligomerization data, we used AlphaFold3 to predict oligomierc states. We first assessed if known oligomeric states are predicted with high confidence scores (ipTM scores). The results established that the predictions matched the experimental SEC-MALS data for all analyzed orthologs (Fig. S2A), on the basis of which we applied the approach to additional orthologs. The mapping of experimentally or predicted quaternary structure showed that the common diDNase/NrnC ancestor A0 was a dimer and the diDNase lineage kept this trait without developing tendencies to assemble into higher oligomeric structures. All extant NrnC orthologs, as well as NrnC ancestors starting from A3, form octamers, suggesting NrnC proteins acquired this trait after ancestor A2 and kept it throughout evolution. This indicated that the ability to form octamers occurred early in evolution, in the transition between A2 and subsequently emerged orthologs. The emergence of octamerization can also be traced by looking at the presence of the primary amino acid sequence elements involved in octameric interfaces based on structural analysis (2) to the same point in ancestry (Fig. S3).

Both NrnC-specific phenotypes, absence of substrate preference towards DNA and octameric arrangement, occurred simultaneously during NrnC specification. Consistent with this conclusion, we also determined that the molecular basis for both phenotypes is provided by overlapping structural elements (Fig. 3C). We analyzed the octamerization contacts between dimeric subunits of available NrnC structures, which we determined for proteins from the Alphaproteobacterial order Hyphomicrobiales (*3, 8*). In addition, we here determined crystal structures of additional Alphaproteobacterial NrnC orthologs from *Inquilinus* (order Rhodospirillales) and *Sphingomonas* (order Sphingomonadales). The octamerization interfaces proved to be conserved across all NrnC ortholog structures (Fig. S2B). As discussed before, the α4-α5 loop in NrnC adopts a particular conformation that contributes to creating more space in the active site of the protein (Fig. 2A-B). At the same time, the loop participates in building interactions between dimeric subunits inside the NrnC octamer (Fig. 3C). Another structural feature implicated in defining substrate promiscuity of NrnC by our structural analysis, W^197^ (Fig. 2B), creates a cation-π bond with R^37^ of the neighboring subunit inside an octameric assembly (Fig. 3C). The conformation of the α4-α5 loop and position of residue 197 in a diDNase are not compatible with NrnC-like inter-subunit contacts - both elements are too far apart to engage in such an interaction (Fig. 3D).

In addition to protein-protein interactions, the α4-α5 loop in NrnC proteins participates in coordination of the 3’ base of the dinucleotide substrate. We have explored the significance of this coordination for NrnC function in the next section.

**Figure S2.**
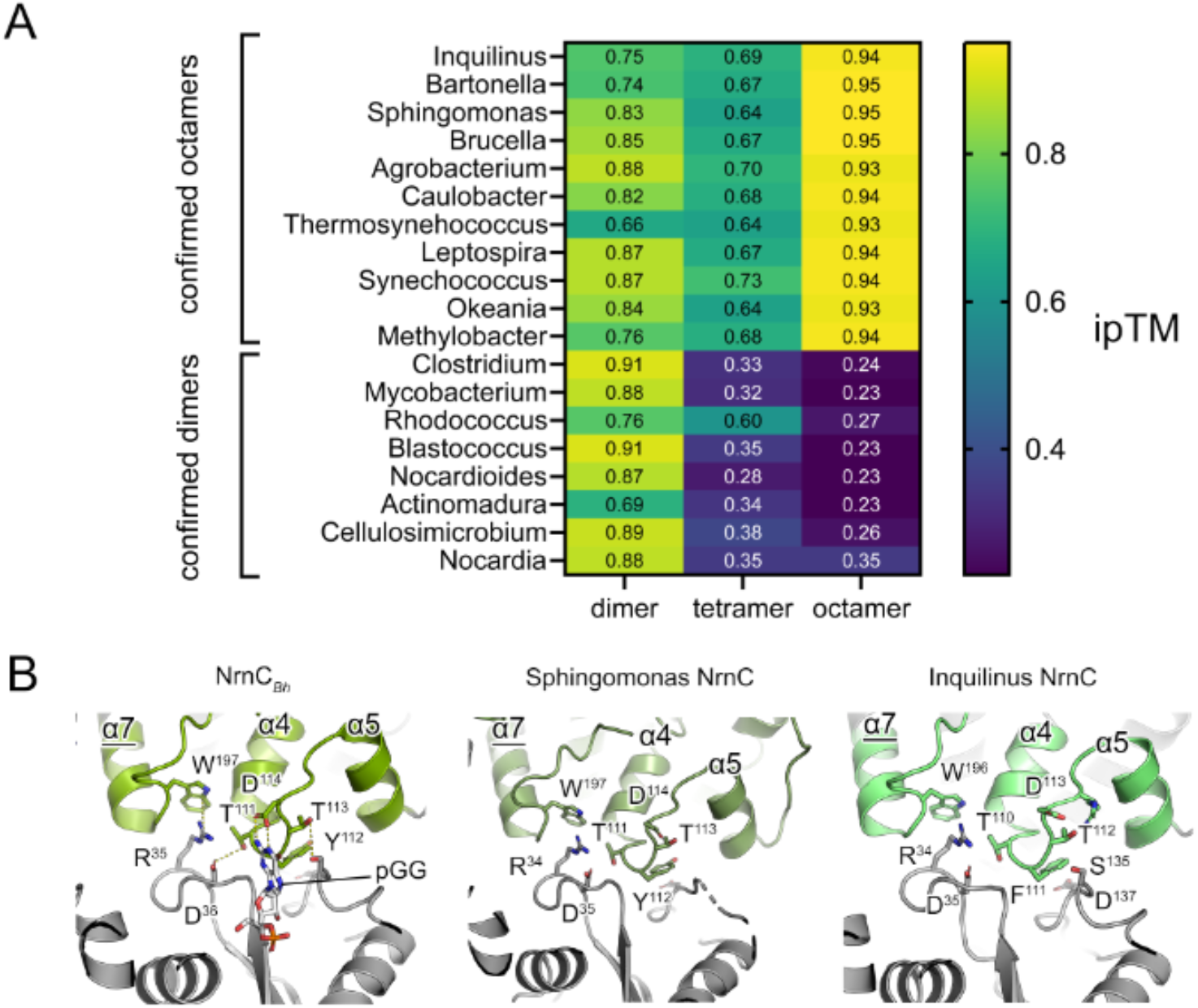
Oligomeric states and octamerization interfaces of diDNases and NrnC orthologs. **A**. AlphaFold3 was used to predict the oligomeric states of NrnC orthologs and diDNases. The ipTM scores of dimeric, tetrameric and octameric states of different proteins are plotted as a heatmap. Actual oligomeric states were confirmed experimentally and are indicated. **B**. Structural comparison of the octamerization interfaces of *Inquilinus, Sphingomonas*, and *Bartonella* NrnC.

### Role of octamerization for NrnC enzymatic activity

Since two NrnC specification events, disappearance of DNA preference and emergence of octamerization, co-occurred, we next investigated whether these characteristics depend on one another or emerged coincidently. In order to answer this question, we tried extensively to create artificially dimeric NrnC mutants for measuring enzyme function by breaking the octameric interface of a NrnC, however, these variants could not be expressed recombinantly. As an alternative, we assessed whether there were natural NrnC orthologs that would disassemble into smaller subunits at lower protein concentrations, allowing us to measure the catalytic activity of their subunits. We identified two alphaproteobacterial NrnC orthologs, from *Caulobacter* and *Sphingomonas*, which displayed instability of octameric arrangements in our SEC-MALS experiments. Both of the orthologs existed as mixtures of dimers, tetramers, and octamers at micromolar concentrations used for the analysis (Fig. 4A). When diluted further to 100 nM concentration, they appeared almost entirely as dimers (>85%) according to mass photometry, an experimental method better suited for mass determination at low analyte concentration (Fig. 4B and Table S4). We utilized the ability of these NrnC orthologs to stay dimeric at low concentrations to measure the enzymatic activities originating from NrnC in its dimeric state. Both *Caulobacter* and *Sphingomonas* NrnC at 100 nM processed RNA and DNA dinucleotides indiscriminately similar to all other NrnC orthologs tested so far (Fig. 4C, Table S2). This result indicates that the absence of DNA preference does not require octamerization of NrnC.

**Figure 4.**
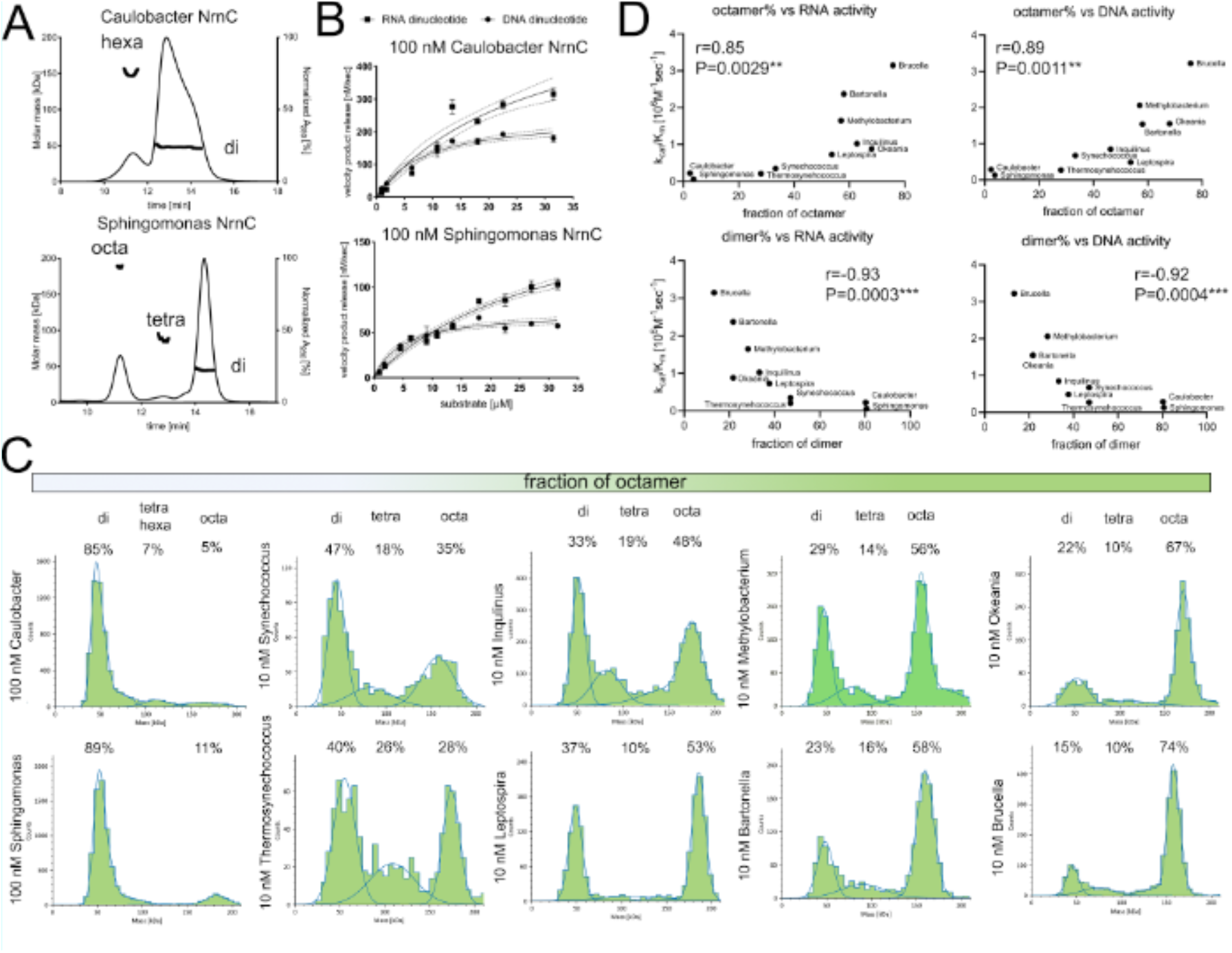
Assessment of NrnC oligomerization and its relation to enzymatic activity. **A**. Oligomeric states of *Caulobacter* and *Sphingomonas* NrnC determined by SEC-MALS at micromolar protein concentrations. The normalized absorbance at 280 nm (chromatogram lines, right axis) and molar mass values (dots across peaks, left axis) were plotted against the SEC elution volume. **B**. Enzymatic activities of *Caulobacter* and *Sphingomonas* NrnC towards DNA and RNA dinucleotides measured at 100 nM when the proteins are >85% dimeric. **C**. Mass Photometry histograms obtained for different NrnC samples at nanomolar concentrations. The mean values of Gaussian fits correspond to the theoretical molecular mass of dimer, tetramer, hexamer, and octamer states of NrnC orthologs. The measurements were done in triplicates and representative histograms are shown. **D**. Correlations between octamer or dimer fractions of 10 different NrnC orthologs and levels of their enzymatic activity. The r factors and P values derived by Spearman nonparametric correlation are indicated.

Since octamerization of *Caulobacter* and *Sphingomonas* NrnC depended noticeably on protein concentration, we next assessed oligomeric states of all other NrnC orthologs at concentrations used previously for determining enzyme kinetics (Table S4, S5). We found that all NrnC orthologs tested existed as mixtures of dimers, tetramers, and octamers at 10 nM with the fraction of octamer varying across the different proteins (Fig. 4B). From this group, the *Brucella* and *Okeania* NrnC orthologs displayed the highest fraction of octamers. We next asked whether there is a link between the level of octamerization and enzyme activity among NrnC orthologs. To this end, we ran correlation tests using the K_cat_/K_m_ of ten NrnC orthologs towards either RNA or DNA dinucleotide and their dimer or octamer fractions (Fig. S3 and Table S2). We found that for both substrates there is a strong positive correlation between enzyme activity and octamer fraction and strong negative correlation between level of activity and dimer fraction (Fig. 4D). Moreover, for the enzyme with the highest octamer fraction, NrnC from *Brucella*, we observed enzyme kinetics that fit a sigmoidal curve, indicating positive cooperativity (Fig. S3). Together, these data indicate that octamerization in NrnC orthologs increases global enzymatic activity without affecting substrate specificity. This observation can be rationalized by the structural insight showing that the α4-α5 loop involved in inter-dimer interactions within octameric assemblies also takes part in coordinating the 3’ base of the dinucleotide substrate (Fig. 3D), which may enhance binding of substrate during catalysis. Considering that the inter-dimer interface residues also make contact to a substrate (Fig. 4C), we asked whether the presence of substrates promotes or stabilizes octameric assemblies. To answer this, we measured oligomeric state distributions of 50 nM of two NrnC orthologs, one with the weakest ability to form octamers when diluted – from *Sphingomonas* – and one with intermediate level of this ability – from *Thermosynechococcus*, in the presence or absence of substrates (Fig. S3C). We did not observe any significant increase in octamer fractions when substrates were present in these experiments, suggesting that oligomer formation and stability do not rely on the substrate.

**Figure S3.**
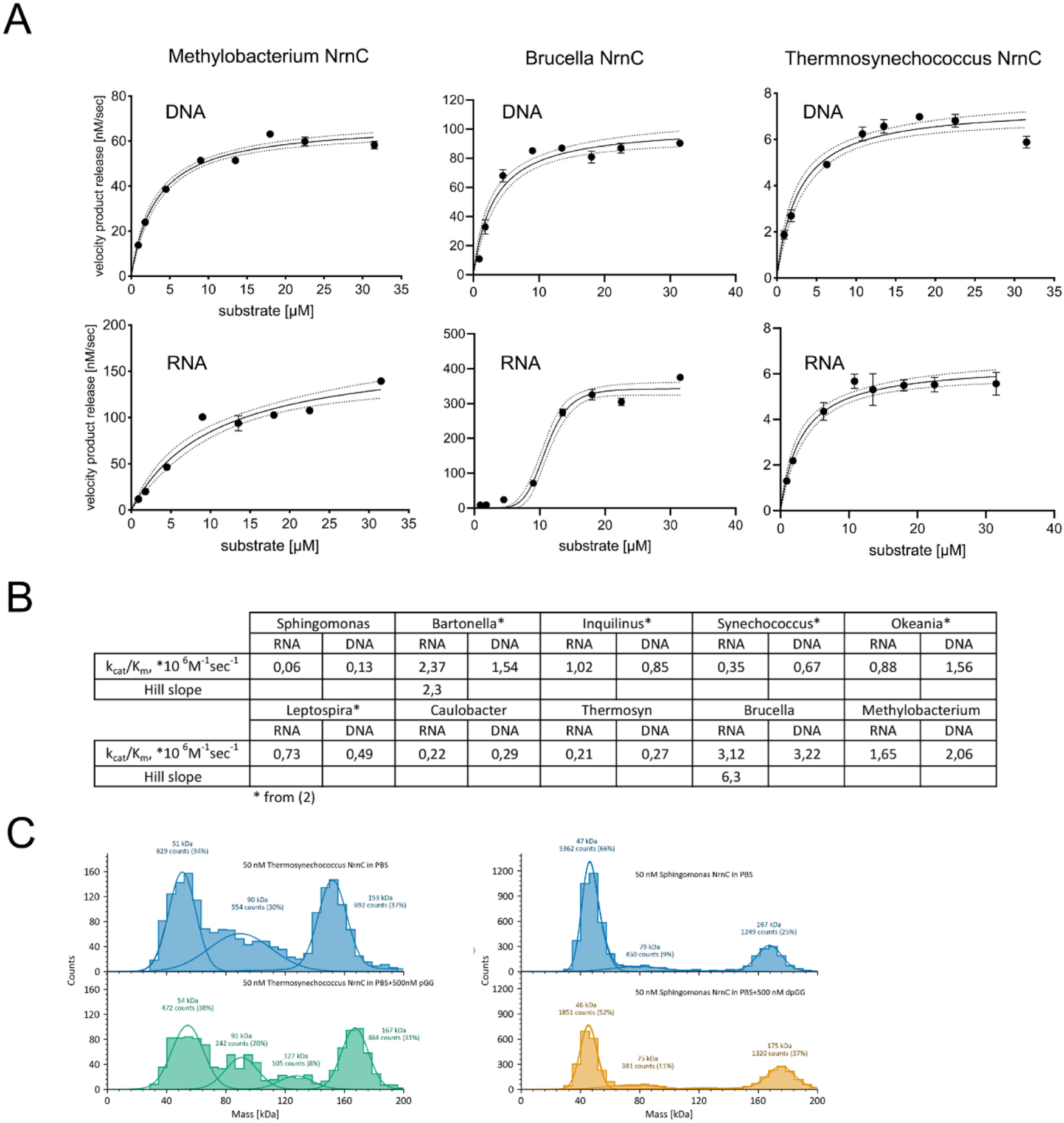
Kinetic parameter and octamerization of NrnC. **A**. Enzymatic activities of three NrnC orthologs towards DNA and RNA dinucleotides. Data points were measured in triplicate and fitted using the Michaelis-Menten or Allosteric Sigmoidal models. Mean values and standard deviations (SDs) are presented. The dotted lines indicate the 95% confidence intervals of model fitting. **B**. Catalytic efficiencies, k_cat_/K_m_, of ten different NrnC orthologs towards RNA and DNA dinucleotides. The efficiency values were used for correlation analysis shown in Figure 4D. **C**. Representative Mass Photometry histograms of *Sphingomonas* and *Thermosynechococcus* at 50 nM of protein concentration in absence and presence of 500 nM dinucleotide substrates.

**Table S2.** Enzyme kinetics parameters of NrnC proteins towards DNA and RNA dinucleotide substrates.

|  | Sphingomonas |  | Caulobacter |  | Thermosyn |  | Brucella |  | Methylobac |  |
| --- | --- | --- | --- | --- | --- | --- | --- | --- | --- | --- |
|  | RNA | DNA | RNA | DNA | RNA | DNA | RNA | DNA | RNA | DNA |
| [E], nM | 100 |  | 100 |  | 10 |  | 10 |  | 10 |  |
| V<sub>max</sub>, nM/sec | 218,3 | 72,08 | 629,1 | 250,5 | 6,452 | 7,456 | 343,1 | 102,7 | 173,7 | 68,23 |
| K<sub>m</sub>, μM | 34,15 | 5,44 | 28,45 | 8,521 | 3,058 | 2,764 | 11,01 | 3,186 | 10,52 | 3,32 |
| k<sub>cat</sub>, sec<sup>-1</sup> | 2,18 | 0,72 | 6,291 | 2,505 | 0,6452 | 0,7456 | 34,31 | 10,27 | 17,37 | 6,823 |
| k<sub>cat</sub>/K<sub>m</sub>,<br>×10<sup>6</sup> M<sup>-1</sup>s<sup>-1</sup> | 0,060 | 0,130 | 0,221 | 0,294 | 0,211 | 0,270 | 3,116 | 3,223 | 1,651 | 2,055 |
| DNA/RNA | 2,167 |  | 1,329 |  | 1,279 |  | 1,034 |  | 1,245 |  |

## Discussion

The evolution of diDNases and NrnC orthologs provides intriguing insights into the adaptive dynamics of nucleases within a shared structural fold. Both of the enzymes evolved independently from an ancient common ancestor with intermediate substrate preference. Additionally, our findings suggest that mobile diDNases have persisted as distinct enzymes in Actinomycetes and Clostridia for as long as house-keeping NrnC orthologs have done. This highlights the importance of diDNase activity to bacterial populations, likely due to their beneficial roles e.g., in phage defense or DNA mobility (*2*). Our results also indicate that diDNases have tolerated a relatively high diversity throughout evolution, likely driven by their faster evolution rate as part of MGEs (*9*), without affecting the conservation of their substrate preference and oligomeric state. The hallmarks of gene mobility in the entire diDNase phylogenic clade, such as heterogenic phylum distribution and high sequence diversity, together with the fact that diDNase split from NrnC orthologs early in evolution suggest diDNases became mobile early on.

Both diDNases and NrnC enzymes appear to have evolved from a low-activity ancestor and progressively developed higher enzymatic activity, with diDNases specializing in DNA and NrnC orthologs developing a broader substrate preference. This suggests that the primary evolutionary driver for both groups was the need for high nuclease activity. At the same time, NrnC-specific evolutionary pressure was to keep both RNase and DNase activities more or less equally high. This result emphasizes the potential importance of dinuclease activity toward DNA for the enzyme’s biological function, making diDNases dispensible under normal growth conditions. Notably, while both bacterial house-keeping dinucleases, NrnC (*2, 3*) and Orn (*2, 10*), as well as functionally related nanoRNases, NrnA (*11*) and NrnB (*12*), show high DNase activity, the physiological significance of this activity has not been fully explored. Organisms that contain a diDNase gene also encode an Orn ortholog with activity against both, DNA and RNA (*2*), suggesting that both enzymes play a role for cellular homeostasis or survival, with diDNases likely providing a more specific, potentially context-dependent function. At the same time, expression of diDNases in a *P. aeruginosa orn* deletion strain yields partial complementation, indicating that the activity of Orn orthologs against DNA dinucleotides likely contributes to the full spectrum of the deletion phenotype (*2*). In this regard, the ancestors of diDNases and NrnC orthologs resemble diDNase, providing only a partial rescue of the *orn* deletion, highlighting the physiologically importance of increased RNase activity in the Orn and NrnC clades (*2*–*4, 13*).

We showed that the propensity of NrnC orthologs to form octamers correlates with increased catalytic activity, suggesting that octamerization could have been selected for in evolutionary history to optimize enzymatic function or robustness. However, octamerization does not appear to be strictly required for the processing of both, RNA and DNA dinucleotides, since NrnC orthologs at low concentration disassemble into their dimeric subunits with comparable substrate preference to the octameric assemblies at higher protein concentration, albeit with reduced catalytic efficiency towards both substrates (Fig. 4). This observation does not rule out that octamerization may have been important in manifesting such a promiscuous substrate profile during evolution. Based on our structural analysis, octamerization requires a certain dimer subunit configuration (Fig. 3). At the same time, we observed that this configuration and its associated localized conformational changes play a crucial role for opening up the active site for RNA dinucleotides (Fig. 2). In contrast, the arrangement of protomers within diDNase dimers is incompatible with octamerization (Fig. 3), and the active sites are optimized for DNA dinucleotides. Hence, it is conceivable that octamerization aided in the rearrangements at the dimer interfaces, rendering NrnC orthologs robust enzymes maintaining high catalytic efficiency towards both RNA and DNA substrates. At low enzyme concentrations, the octamers break apart but the particular dimer subunit configuration likely persists, explaining the unaltered substrate specificity. Loosing contacts at the inter-dimer interface with the 3’ base of the substrate upon octamer disassembly may explain the reduced overall activity under these conditions (Figs. 3 and S2).

Enzymes performing analogous function to NrnC in other bacteria, namely Orn, NrnA and NrnB, are dimeric and do not form higher assemblies, however, the level of their nuclease activity is comparable with that of extant NrnC orthologs (*2, 4, 11, 12*). Hence, high nuclease activity can be achieved without large oligomeric forms and NrnC-specific octamerization may have other roles for the function of the enzyme, such as interactions with cellular factors, allosteric regulation, or as an additional size filter considering that the active sites line the walls of a narrow channel of the octameric assembly. Few recent reports on protein oligomerization suggested that the formation of high oligomers in metabolic enzymes (*14*) or other proteins (*15*) is generally a neutral evolutionary event that does not provide benefits. Our data provides evidence that, in the case of NrnC, octamerization is beneficial for achieving high enzymatic activity.

## Materials and Methods

### Protein expression and purification

Genes encoding ancestral proteins, codon-optimized for expression in *E. coli*, were synthesized and fused with a His_6_-tagged small ubiquitin-like modifier (SUMO) at the N-terminus in a modified pET28a vector (GenScript, Novagen). The resulting constructs were transformed into One Shot BL21(DE3) cells (Invitrogen). Transformed cells were grown in lysogeny broth (LB) supplemented with 50 μg/ml kanamycin overnight at 37°C with shaking. Auto-induction media ZYM-5052 (*16*) supplemented with 100 μg/ml kanamycin was inoculated at a 1/100 ratio with the overnight culture, grown at 37°C with shaking for 2 h, and then at 18°C with shaking for 18-20 h. Cells were harvested by centrifugation and the pellets were stored at −70°C until purification.

For purification, cell pellets were thawed and resuspended in lysis buffer (50 mM Tris-HCl, 500 mM NaCl, 25 mM imidazole, 1 mM PMSF, 3 mM β-mercaptoethanol [pH 8.5]). Cells were lysed by sonication, and the lysates were clarified from the insoluble material by centrifugation. Clarified lysates were incubated with Ni-NTA resin (Qiagen) with gentle agitation at 8°C for 1 h. The resin was collected by centrifugation and washed 3 times with five resin volumes of lysis buffer, and placed in a gravity column. His_6_-tagged Ulp1 protease was added to the NiNTA resin in 1-2 resin volumes of lysis buffer, the column was sealed, and the resin was incubated at 8°C for 1 h with rotation to allow for cleavage of the His_6_-SUMO tag. Subsequently, the flow through containing the untagged protein of interest was collected and concentrated on centrifugal filter units (Amicon Ultra-4, 10 kDa cut-off). EDTA (pH 8.5, final concentration of 10 mM) was added to the concentrated protein to chelate divalent cations and further purified by gel filtration using either HiLoad 16/600 Superdex 200 pg or Superdex 200 Increase 10/300 GL columns (Cytiva) equilibrated with gel-filtration buffer (for ancestor A0, 25 mM HEPES-NaOH and 500 mM NaCl [pH 7.5]; for all other, 25 mM HEPES-NaOH and 150 mM NaCl [pH 7.5]). Protein-containing fractions were pooled, concentrated, flash-frozen in liquid nitrogen, and stored at −70°C.

### Size-exclusion chromatography-coupled multiangle light scattering (SEC-MALS)

Purified proteins at 1-2 mg/ml (~40-80 μM) were injected onto a Superdex 200 Increase 10/300 GL column (Cytiva) equilibrated with gel-filtration buffer at room temperature. The gel-filtration setup was coupled inline to a static multi-angle light scattering detector (miniDAWN, Wyatt Technology, Waters) and a refractive index detector (Optilab T-rEX, Wyatt Technology, Waters). Data analysis was performed using the Astra software (version 8.0.2.5, Wyatt Technology, Waters), yielding molar mass values across elution peaks of samples, as well as weight fractions of each molar mass species.

### Enzyme kinetics

Substrate hydrolysis by NrnC homologs was measured by fluorescence unquenching of nucleotides containing 2-aminopurine as one of the bases (*17*). All substrates were purchased 5’-phosphorylated from Dharmacon. Enzymatic activity was measured as the rate of increase in the fluorescence intensity of cleaved 2-aminopurine with excitation at 310 nm and emission at 375 nm over a series of initial substrate concentrations (0.9-45 μM). Enzyme concentrations ranged from 10 to 500 nM depending on level of activity and are indicated on particular results. The reactions were assembled by mixing reaction buffer (20 mM Tris-HCl, 150 mM KCl [pH 7.9]) with a substrate stock made in water and an enzyme diluted with dilution buffer (20 mM Tris-HCl, 150 mM KCl, 4 mM EDTA [pH 7.9]). 45 μL of the resulting mixes were placed in wells of a 96-well plate and reactions were started by adding 5 μL of start buffer (20 mM Tris-HCl, 150 mM KCl 100 mM MgCl2 [pH 7.9]). The fluorescence intensity was recorded in triplicate every 10 s for 3 min at 21°C using a Tecan Infinite M Plex plate reader. The initial slopes of the resulting time-course data were determined and converted into the velocities of product release using a calibration curve. Calibration curves were created for each type of substrate tested by incubating reactions for 5-10 hours in a sealed plate to allow for compete substrate conversion and plotting final fluorescence intensity values against initial substrate concentrations. The velocities of product release were plotted against initial substrate concentrations and fitted with enzyme kinetics models in GraphPad Prism 10.

### Crystallization and determination of structures

All crystals were obtained in sitting-drop setups at 19°C. *Sphingomonas* NrnC was crystallized in 0.005M Cobalt (II) chloride hexahydrate, Nickel (II) chloride hexahydrate, cadmium chloride hydrate, Magnesium chloride hexahydrate, 0.1M HEPES (pH 7.5), 12% (w/v) PEG 3350. *Inquilinus* NrnC was crystallized in 0.2 M magnesium chloride, 0.1 M Bis-Tris (pH 5.5), 25% PEG 3350. Ancestor A1 was crystallized in 0.1M MES (pH 5), 20% (w/v) PEG 6000. The crystals were cryoprotected by either xylitol or a low-viscosity oil prior to flash cooling in liquid nitrogen. Diffraction data were collected at beamline P11 at the Deutsches Elektronen-Synchrotron (DESY) and processed using XDS (*18*). Phase information was obtained by molecular replacement using Phaser (*19*). The models were iteratively refined using Phenix.refine (*20*) and manually built in Coot (*21*). Structural figures were prepared using PyMOL (version 2.5.0; Schrödinger). Data processing and model refinement statistics are presented in Table S3.

**Table S3.** X-ray diffraction data processing and refinement statistics.

| Structure | Ancestor A1 | Sphingomonas NrnC | Inquilinus NrnC |
| --- | --- | --- | --- |
| Resolution range | 43.43 - 1.45 (1.502 - 1.45) | 41.52 - 3.0 (3.107 - 3.0) | 43.14 - 1.75 (1.813 - 1.75) |
| Space group | C 1 2 1 | P 4 2 2 | P 4 2 1 2 |
| Unit cell | 70.903 70.948<br>48.241 90 115.807<br>90 | 103.398 103.398<br>139.393 90 90 90 | 120.708 120.708<br>71.764 90 90 90 |
| Total reflections | 256038 (25793) | 201061 (20544) | 994274 (99824) |
| Unique reflections | 36862 (3622) | 7851 (765) | 53803 (5247) |
| Multiplicity | 6.9 (7.1) | 25.6 (26.9) | 18.5 (19.0) |
| Completeness (%) | 96.65 (95.28) | 99.42 (100.00) | 99.77 (99.54) |
| Mean I/sigma(I) | 14.42 (2.79) | 37.23 (5.29) | 18.30 (1.66) |
| R-merge | 0.06764 (0.7653) | 0.06462 (0.7159) | 0.1888 (3.314) |
| CC1/2 | 0.998 (0.91) | 1 (0.956) | 0.998 (0.689) |
| CC* | 1 (0.976) | 1 (0.989) | 0.999 (0.903) |
| Reflections used in refinement | 36840 (3617) | 7852 (765) | 53778 (5240) |
| Reflections used for R-free | 1600 (157) | 788 (75) | 862 (84) |
| R-work | 0.1666 (0.2239) | 0.2048 (0.4292) | 0.1823 (0.3521) |
| R-free | 0.2023 (0.2851) | 0.2571 (0.5167) | 0.1975 (0.3995) |
| CC(work) | 0.964 (0.938) | 0.940 (0.786) | 0.958 (0.682) |
| CC(free) | 0.947 (0.893) | 0.885 (0.680) | 0.979 (0.639) |
| Number of non-hydrogen atoms | 1802 | 1637 | 3781 |
| macromolecules | 1649 | 1630 | 3165 |
| ligands | 1 | 7 | 6 |
| solvent | 152 | 0 | 610 |
| Protein residues | 202 | 203 | 406 |
| RMS(bonds) | 0.006 | 0.010 | 0.008 |
| RMS(angles) | 0.91 | 1.42 | 1.01 |
| Ramachandran favored (%) | 98.00 | 88.94 | 97.51 |
| Ramachandran allowed (%) | 2.00 | 11.06 | 2.49 |
| Ramachandran outliers (%) | 0.00 | 0.00 | 0.00 |
| Rotamer outliers (%) | 0.54 | 0.00 | 0.60 |
| Clashscore | 3.01 | 7.52 | 4.41 |
| Average B-factor | 30.36 | 106.34 | 26.34 |

### *P. aeruginosa Δorn* complementation assay

Genes of interest were subcloned into a modified pJN105 vector (*22*) for the expression of N-terminal StrepII-tagged proteins in *P. aeruginosa Δorn*. Plasmids were introduced into *P. aeruginosa* by electroporation. Briefly, cells from an overnight culture grown in lysogeny broth (LB) were collected and washed three times with 1 mM MgCl_2_, followed by the addition of plasmid DNA and electroporation. Cells were recovered in 1 ml of LB with shaking at 250 rpm for 1-2 hr at 37 °C and plated on LB agar containing 60 μg/ml gentamicin. For complementation assays, fresh LB supplemented with 60 μg/ml gentamicin was inoculated with cells harboring a plasmid, followed by overnight growth. The resulting cultures were diluted to optical density at 600 nm (OD_600_) of 0,1, followed by preparation of serial dilution series. Each dilution (12 μl) was applied to LB agar plates containing 60 μg/ml of gentamicin and 0.2% arabinose. The plates were inverted, allowing the culture to drip down the length of the plate. The plates were incubated overnight at 37°C. Plates were imaged and colony areas of well-separated colonies were determined using particle analysis in ImageJ (*23*). The area of each colony was divided by the average area of wild-type *P. aeruginosa* colonies grown on the same plate to allow for normalization across all plates. A minimum of 50 colonies were quantified.

### Western blotting

*P. aeruginosa* expressing StrepII-tagged ancestors from a plasmid were cultured in LB supplemented with 60 μg/ml gentamicin and 0.2% arabinose overnight at 37°C. Cells (1-3 ml) of each culture were collected and resuspended in SDS-PAGE loading buffer. The samples were incubated at 100°C for 10 min and resolved on 15% SDS-PAGE gels. Proteins were transferred onto a 0.2 μm PVDF membrane using a Trans-Blot Turbo Transfer System (Bio-Rad). The membranes were blocked with EveryBlot Blocking Buffer (Bio-Rad) for 1 h at 20°C, followed by incubation with StrepTag II Antibody-HRP Conjugates (Millipore, Merck, at a 1:10000 dilution) for 1 hour at 20°C. Membranes were washed 3 times for 5 min with PBST and imaged with Amersham ECL Select Western Blotting Detection Reagent (Cytiva) for chemiluminescence detection.

### Mass Photometry

Diluted protein (12 ul, 20-200 nM) were mixed with an equal volume of PBS directly prior to the measurement. Data were recorded at 100 frames per second for 60 s using AcquireMP (Refeyn Ltd, v1.2.1) on Two MP instrument (Refeyn Ltd) and analyzed in DiscoverMP (Refeyn Ltd, v2022 R1). Each protein was measured at least in trice.

### AlphaFold3 modelling of oligomeric states

The oligomeric structures were modelled using AlphaFold3 (*24*) on the AlphaFold Server with default settings and the reported ipTM score were reported.

### Phylogenetic Reconstruction and Ancestral Sequence Reconstruction

Protein sequences from the representative genomes of the Genome Taxonomy Database (release 212) (*25*) were annotated using eggNOG-mapper (version 2.1.3) (*26*) and InterProScan (version 5.64-96.0) (*27*). Sequences annotated as belonging to COG0349 (Ribonuclease D) or having annotated PF01612 domain (3’-5’ exonuclease) were selected and clustered at 60% identity using CD-HIT (version 4.8.1) (*28*). The sequence dataset was curated to remove exceptionally long or short sequences (less than 150 amino acids or greater than 400 amino acids) based on the lengths of known NrnC and diDNase proteins. The remaining sequences were supplemented with 44 sequences from experimentally characterized NrnC and diDNase proteins to give a total dataset of 2,401 NrnC, diDNase, and related nucleases.

The sequences were aligned using Clustal Omega (version 1.2.4) (*29*) and the alignment was trimmed using the ‘gappyout’ method in trimAL (version 1.5.0) (*30*). A phylogenetic tree was constructed using IQ-TREE (version 2.3.5) (*31*) using the best-fit model (LG amino acid exchange rate matrix, empirical amino acid frequencies, with the FreeRate model of rate heterogeneity) and 10,000 bootstraps. The phylogenetic tree was manually rooted based on the positioning of the experimentally characterized NrnC and diDNase proteins, to separate the clads of interest from other related nuclease sequences. The ancestral sequence reconstruction was performed based on this reconstructed tree with GRASP with default settings (version 1.0) (*30*). The phylogenetic reconstruction was visualized using iTOL (*31*).

## Acknowledgments

We acknowledge technical support from the SPC facility at EMBL Hamburg. We further acknowledge DESY (Hamburg, Germany), a member of the Helmholtz Association HGF, for the provision of experimental facilities. Parts of this research were carried out at PETRA III, and we would like to thank Johanna Hakapää and the entire P11 staff for their assistance in using beamline P11. This work was supported by the National Institutes of Health (R01 AI142400 to H.S.). Access to the high-throughput crystallization and biophysical characterization facilities were supported by iNEXT Discovery platform and MOSBRI (funding from the European Union’s Horizon 2020 research and innovation program [101004806 to H.S. and S.M.]). This research was supported in part by the Division of Intramural Research (DIR) of the National Library of Medicine (NLM), National Institutes of Health.” and “This work utilized the computational resources of the NIH HPC Biowulf cluster (https://hpc.nih.gov).

## Competing interests

Authors declare that they have no competing interests.

## Author contributions

Conceptualization: SM, AB, KDT, XJ, HS

Methodology: SM, AB, KDT, AEL

Investigation: SM, AB, KDT, AEL

Visualization: SM, KDT

Supervision: SM, XJ, HS

Writing—original draft: SM, KDT

Writing—review & editing: SM, KDT, XJ, HS

## Notes

### Competing Interest Statement

The authors have declared no competing interest.

